# Rapid Generation of Medical Countermeasure Candidates Via Computational Variation Analysis

**DOI:** 10.1101/604256

**Authors:** Darrell O. Ricke

## Abstract

Rapid responses to emerging infectious diseases are needed for viral and bacterial pathogens. For some pathogens, no medical countermeasures (MCMs) yet exist. Pathogen heterogeneity and antigenic variation lead to immune response escape mutations for some pathogens (e.g., influenza) limiting the effectiveness of medical countermeasures. High throughput sequencing enables characterization of large numbers of pathogen isolates to which residue variation analysis can be applied to identify low variability targets. Multiple approaches are proposed that leverage these low variability targets as the first step of medical countermeasure development. Classes of MCMs informed by this approach include the following: DNA or RNA vaccines, both B-cell and T-cell vaccination strategies, anti-viral RNA targeting, antibody therapeutics, and aptamer targeting of viral protein complex interfaces as potential treatment strategies for infected individuals. Variation analysis-designed countermeasures targeting the Ebola glycoprotein are presented to illustrate the concepts for the proposed multiple targeted countermeasures.

## Introduction

Emerging infectious diseases (EIDs)[1] are emerging at a rate that has not been seen before with SARS, MERS, Ebola, chikungunya, avian flu, swine flu, Zika, and others discovered since the 1970s[2]. Natural sequence variations in pathogens (heterogeneity) can limit the effectiveness of medical countermeasures. Antigenic variation enables some pathogens to avoid targeted host immune responses. For example, pathogens use easily mutable highly antigenic residues to create immune response escape mutant progeny. An ideal universal vaccine would be effective against all strains of a pathogen. Some vaccines are multi-component with multiple selected antigens to provide coverage of multiple pathogen strains. Broadly neutralizing antibodies (bnAb) do not target easily mutated highly antigenic residues. BnAbs have been observed to occur naturally for influenza[3] and HIV[4]; but no methods exist for designing bnAb targets *de novo*. Advances in high throughput sequencing (HTS) technologies are now enabling the rapid sequencing of pathogen genomes isolated from patients and animals. Reverse vaccinology or vaccinomics [5-7] leverages genome-based “omics” tools for vaccine development targeting surface-exposed proteins. A cornerstone in vaccine design is the optimal selection of antigen(s) [8]. Structural vaccinology leverages available protein structural information in the rational design of vaccines. By combining sequence data from different isolates to characterize genomic and protein sequence diversity with protein structural information, the relative variability of individual pathogen protein residues can be estimated and used to create a variability ‘map’. Exposed pathogen surface residues with no or little variability are candidates for bnAb targets (i.e., epitope-guided vaccine design): they are maximally accessible to antibodies and represent a relatively constant target across multiple pathogen strains. In addition to enabling selection of optimal targets for bnAb development, this analytical approach can improve development of other medical countermeasures, such as designing DNA and RNA vaccines, identifying linear peptide segments with no or little variability, which could be leveraged as targets to (1) stimulate T-cell vaccination responses or complimentary RNA (synthetic miRNA or siRNA) targeting of the pathogen genome or RNA molecules. In addition, low variability surface areas offer candidate antibody targets and also aptamer targets. Efficacy studies in animals can be leveraged with FDA Animal Rule (21 CFR 314.601.90) for regulatory review of biological countermeasures against pathogens as an alternative to testing in humans. These multiple targeted strategies can be pursued in parallel for rapidly developing countermeasures for EIDs, including naturally-occurring pathogens that have yet to be discovered and also engineered pathogens.

Ebola, an emerging infectious disease for which multiple outbreaks have occurred, is an example of a known EID that has resisting development of broadly-effective medical countermeasures. The 2014 West African epidemic included 28,610 reported cases with 11,308 deaths[9]. The VSB-EBOV vaccine has been found to be 70 to 100% effective against the Ebola virus but half of the people given the vaccine experience mild to moderate adverse effects that include headache, fatigue, and muscle pain[10]. A monoclonal antibody, MAb114, has been isolated from a human survivor of the 1995 Ebola outbreak in Kikwit[11]. Variation analysis of available Ebola glycoprotein sequences was undertaken for designing alternatives countermeasures. Deadly pathogen outbreaks like Ebola require the rapid development of medical countermeasures; parallel strategies are proposed leveraging low variability pathogen residues.

## Methods

### Rational Targeting of Pathogens

Optimal targets for countermeasure development can be determined by combining available protein sequences (reverse vaccinology) with protein structure information (structural vaccinology) to identify targets with low variability for developing multiple broadly neutralizing countermeasures (e.g., epitope-guided vaccine design). Low variability linear protein peptides and clustered surface residues can be targeted with multiple strategies depicted in Figure 1. First, pathogen genes can be used with both DNA and mRNA subcomponent vaccination strategies. Second, multiple low variability linear peptides can be linked in a vaccination strategy to stimulate T-cell immune responses to the target pathogen peptides; these target peptides can be linked together to leverage the nucleic acid vaccination approach. Third, both linear and non-linear low variability surface residues can be leveraged to develop broadly neutralizing antibodies. Fourth, low variability protein surface residues that are buried can be targeted with aptamers to block virus particle protein complex assemblies within infected cells. Fifth, virus RNA molecules can be targeted with complimentary synthetic miRNA or siRNA molecules to elicit cellular double stranded RNA anti-virus responses.

**Figure 1.**
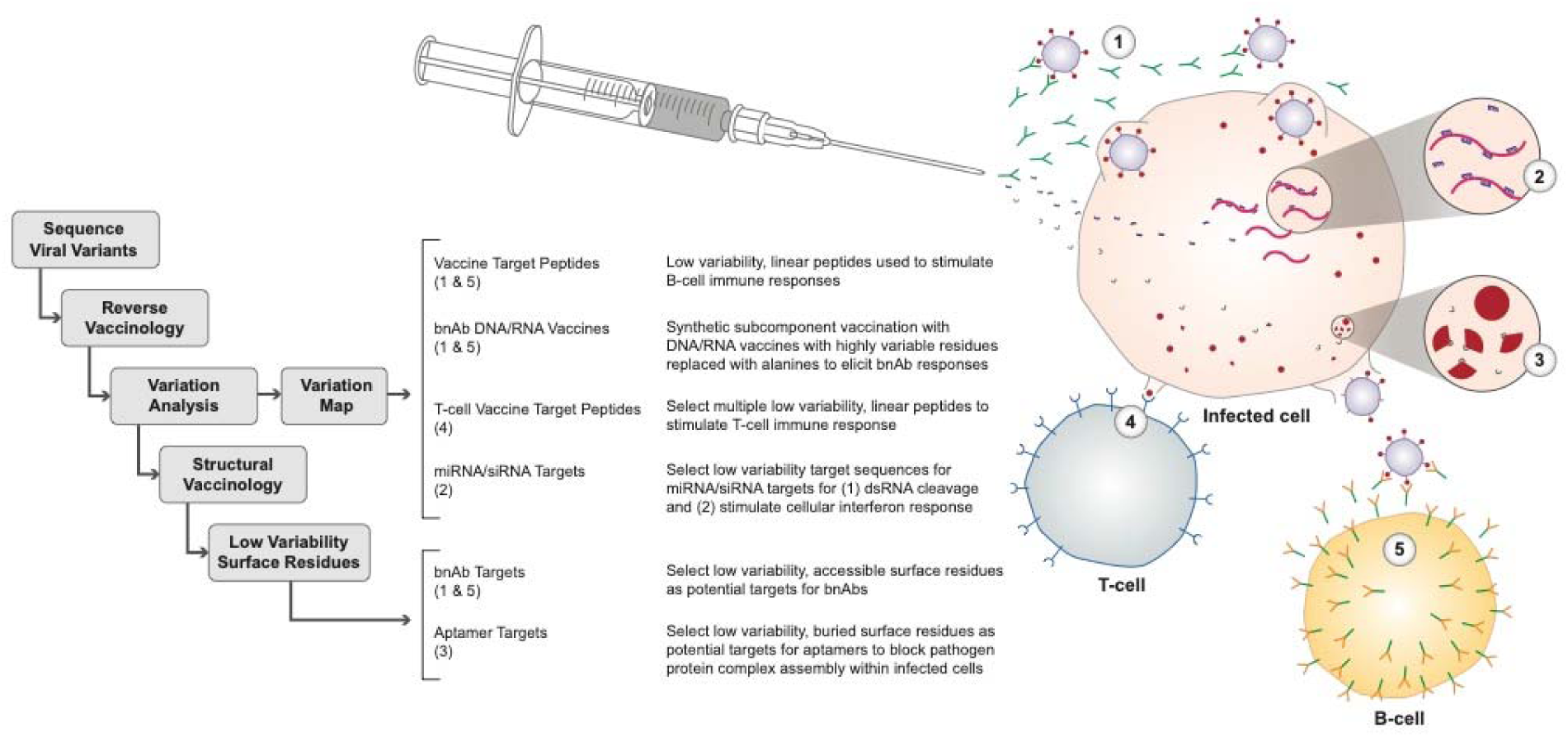
Rational countermeasure design strategies and notional diagram: (1) antibody therapeutic, (2) anti-viral RNA to invoke interferon response, (3) aptamer targeting of viral complex surface interfaces, (4) vaccination for T-cell immune responses, and (5) vaccination for B-cell antibody responses.

These rational countermeasure design strategies can be developed and tested in parallel. Combinations of these approaches may provide enhanced responses (i.e., synergy) than individual treatment responses for either vaccination or treatment modalities. The countermeasures can be developed and optionally evaluated in *in vitro* efficacy assays when time permits. If rapid responses are needed, these countermeasure strategies can be evaluated in animals. Isolation of monoclonal antibodies (mAbs) is possible for the subcomponent and targeted low variability surface epitopes approaches.

### Identification of Low Variability Surface Epitopes

Available pathogen protein sequences can be aligned with a high quality multiple sequence alignment tool (i.e., Dawn[12] or Clustal-Omega[13]). For pathogens with limited available data, sequences and structure data from evolutionarily related proteins can be leveraged. Variation analysis of each alignment position provides a map of known sequence variants; this can be directly leveraged to identify low variability linear peptides with little or no variation. Variation analysis can be combined with protein structure information to identify low variability surface clusters for rational countermeasure targeting.

A total of 4,433 Ebola glycoprotein sequences were selected from GenBank[14] and aligned and variation characterized with Dawn[12]. The Ebola glycoprotein 5KEL[15] structures was selected from PDB[16, 17] for visualizing variation results. Dawn variation results were visualized with the Jmol[18] protein structure viewer. Variance analysis for the Ebola proteins GP, L, NP, VP24, VP30, VP35, and VP40 are provided in a supplemental Excel file.

### Subcomponent Targeting

Pathogen subcomponents can used to illicit immune responses as part of a vaccination strategy. Delivery approaches include both DNA and RNA vaccines. *In vitro* transcribed (IVT) mRNA and self-amplifying mRNA (SAM) vaccines based on an alphavirus genome have been developed[1]. These nucleic acid approaches can include target pathogen genes or customized proteins to stimulate antibody or T-cell immune responses.

### Broadly Neutralizing Antibody Development

Broadly neutralizing antibodies can be developed by selecting antibodies that target low variability pathogen surface residues. If the residues are continuous in a linear peptide, that peptide could be used to elicit antibody responses in animals. Alternatively, highly variable residues pathogen residues can be replaced with alanine residues and the synthetic proteins with these modified protein sequences used to elicit antibody responses in animals. Antibodies to low variability pathogen residues should have the potential to be effective against multiple strains of the pathogen that all share these low variability surface residues. Broadly neutralizing antibodies can be used as passive immunization treatment to exposed individuals.

### Targeting Low Variability Peptides to Elicit T-cell Immune Response

Variation analysis can identify multiple linear peptides with limited or no variation; synthetic proteins containing multiple of these peptide targets can be used to stimulate T-cell immune responses as part of a vaccination strategy. Target peptides can be linked together for simultaneous stimulation for multiple peptide targets. Nucleic acid vaccine approaches can be leveraged to deliver these low variability peptides combined together into synthetic protein(s).

### Targeting virus RNA molecules with small complimentary RNA molecules (miRNA/siRNA)

The linear low variability pathogen epitopes identified can be leveraged as RNA targets for synthetic complimentary RNA targeting some viral RNA genomes and expressed viral RNAs to elicit interferon (IFN) system response to double stranded RNA [19] and cellular anti-viral defense double stranded RNA cleavage.

### Aptamer Targeting of Low Variability Pathogen Protein Complex Interfaces

The interfaces between pathogen protein complexes are potential targets for blocking complex assembly. Frequently, pathogen proteins bind to themselves in multimers or with other pathogen molecules; low variability surface epitopes at these protein complex interfaces will not be exposed on the pathogen protein surface. Aptamers are oligonucleotide or peptide molecules that bind to a specific target molecule. Targeted aptamers binding to these low variability surface epitopes may block assembly of pathogen subcomponents to interfere with assembly of progeny pathogen protein complexes.

### Leveraging Immune Epitope Database

Known epitopes for pathogens are collected in the Immune Epitope Database (IEDB)[20]. Known epitopes can be compared with epitopes identified by variation analysis, to identify available reagents and overlaps between predicted targets and known epitopes.

## Results

All available Ebola glycoprotein (GP) sequences were downloaded from GenBank[14]. A multiple sequence alignment of these sequences was created using Dawn[12]. This alignment is illustrated in Figure 2 with the addition of the Marburg virus glycoprotein. The GP variability observed across Ebola isolates is plotted in Figure 3 and candidate low variability targets details are in Table 1; the best linear peptides with low variability can also serve as complimentary RNA target epitopes (Table 2). The best overlaps of target epitopes with known epitopes is included in Table 1 with variable residues positions highlighted with a light blue font color. Three low variability targets occur at the glycoprotein trimer interface (Figures 4) and two at the base of the glycoprotein at the potential anchoring site with the viral envelope (Figures 5 & 6). A non-linear surface target in the glycan cap is illustrated in Figure 7.

**Table 1.**
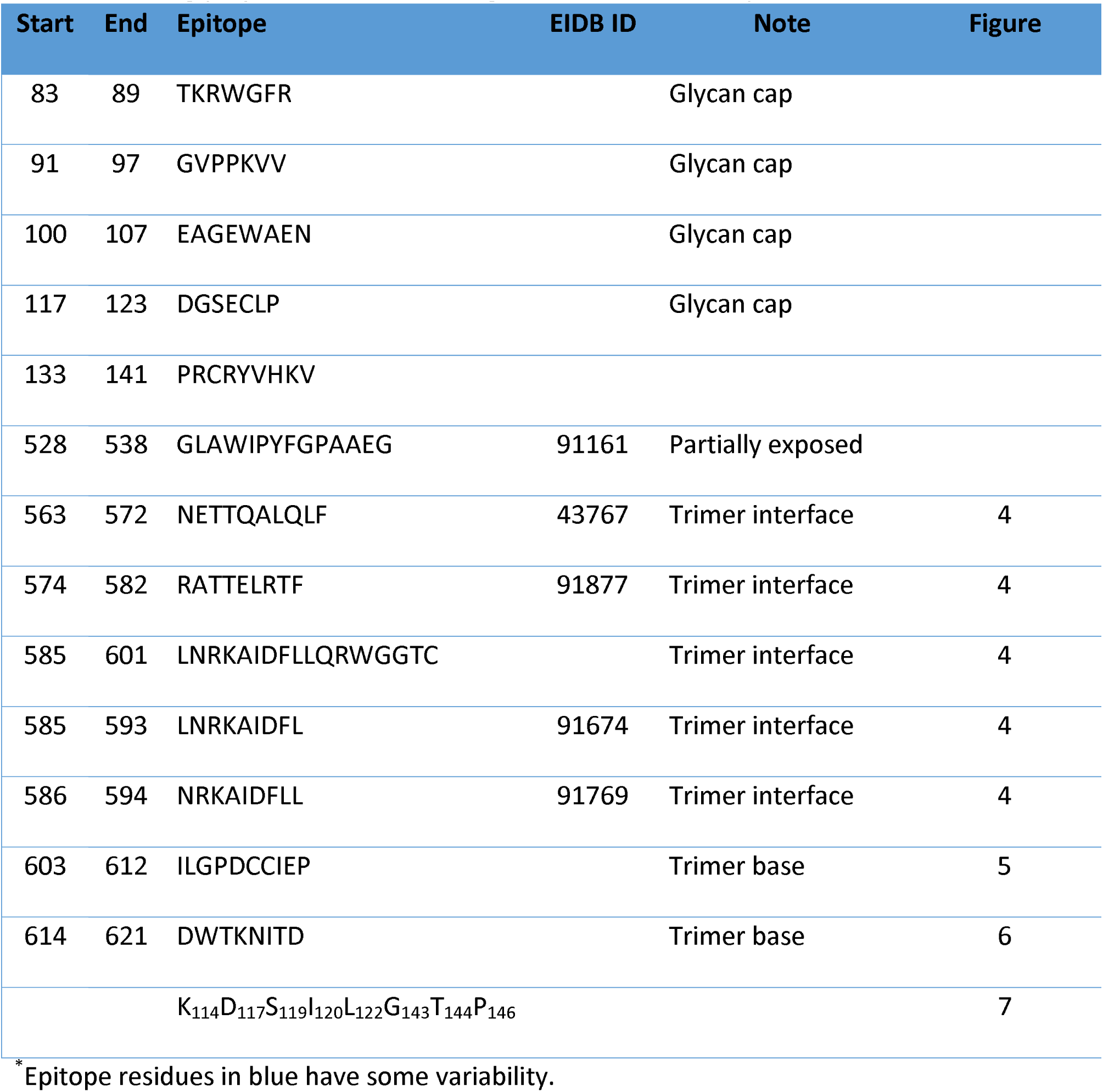
Ebola glycoprotein candidate targets with low variability

**Table 2.**
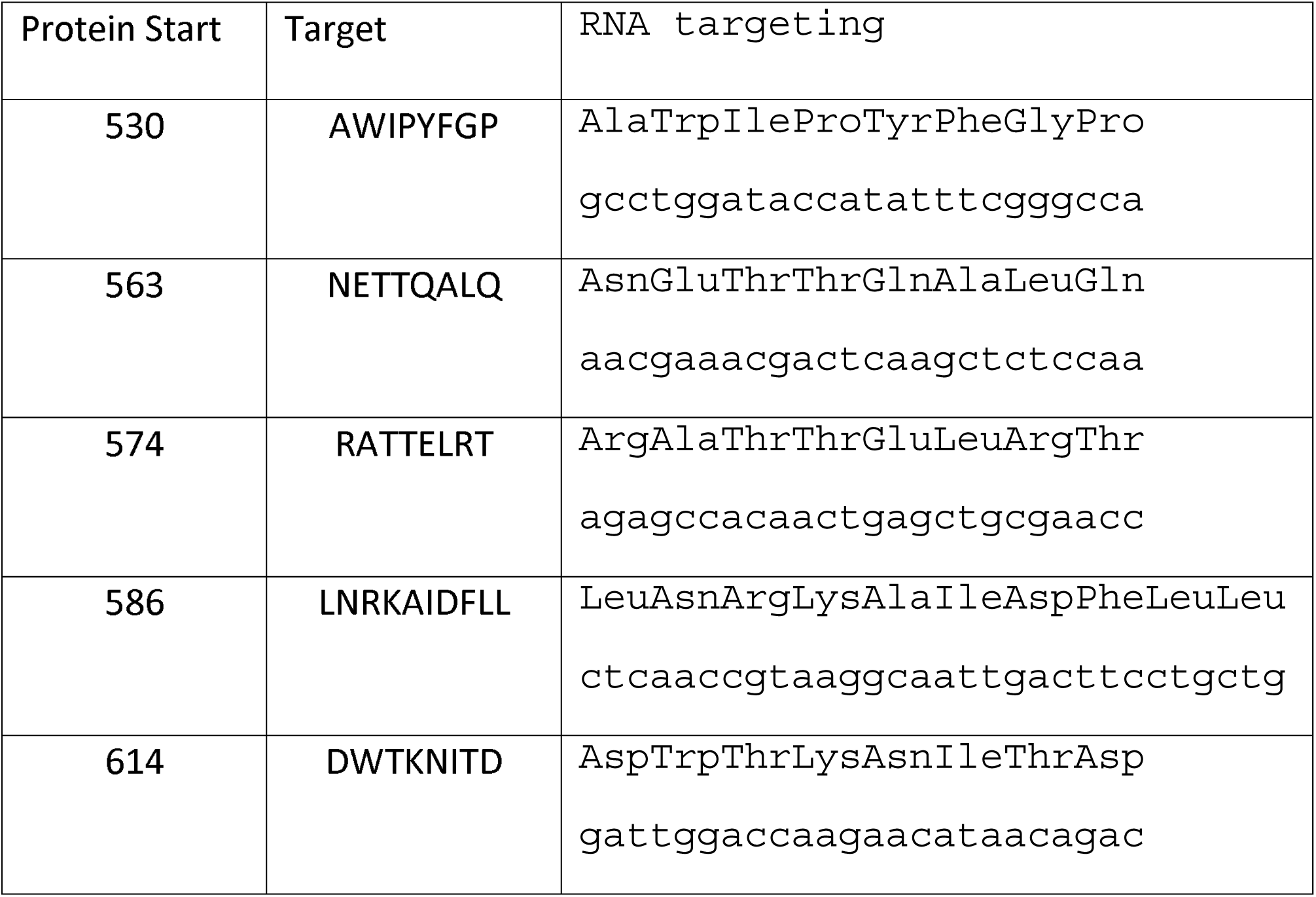
RNA targets for Ebola glycoprotein candidate peptide targets

**Figure 2.**
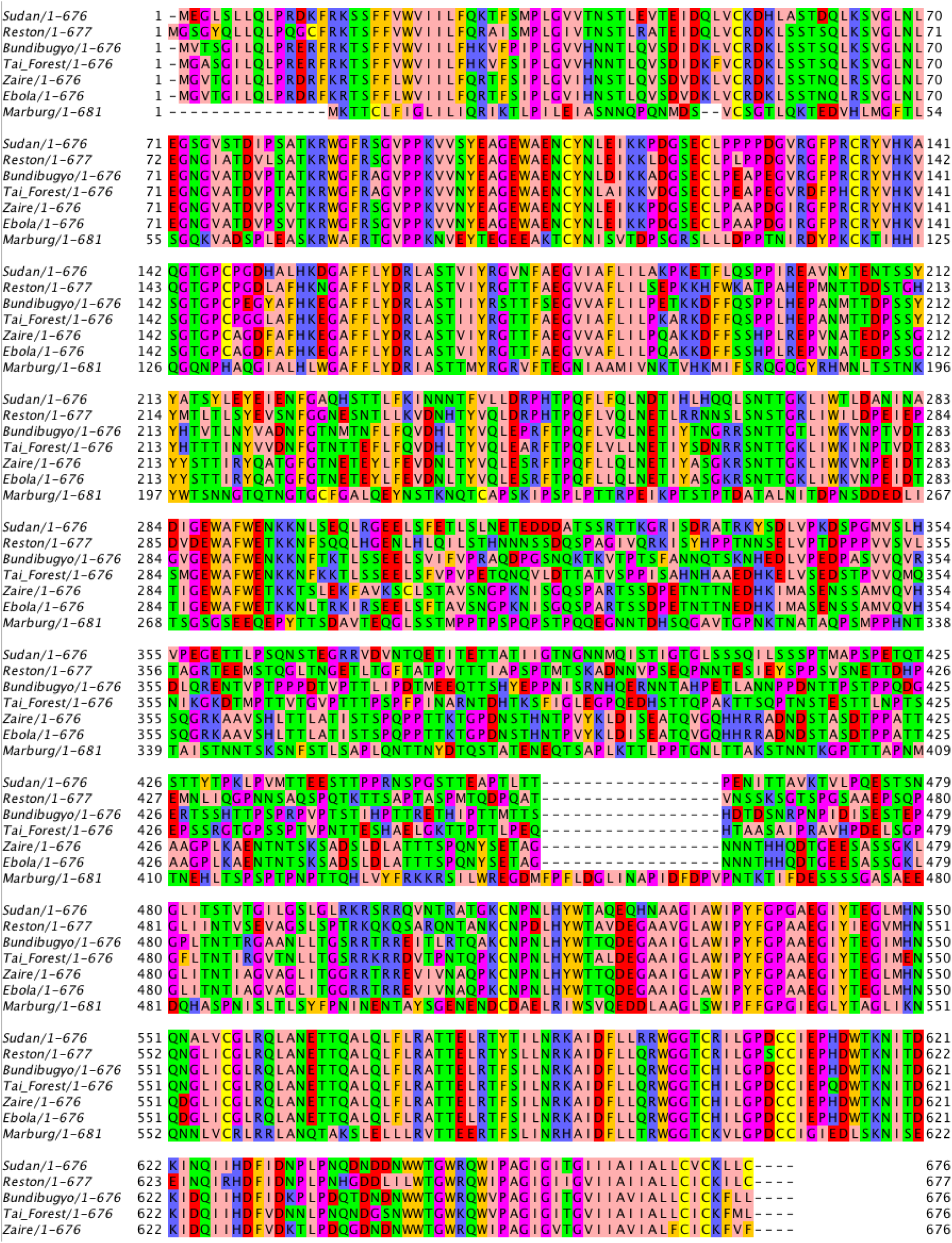
Ebola and marburg viruses glycoprotein alignment

**Figure 3.**
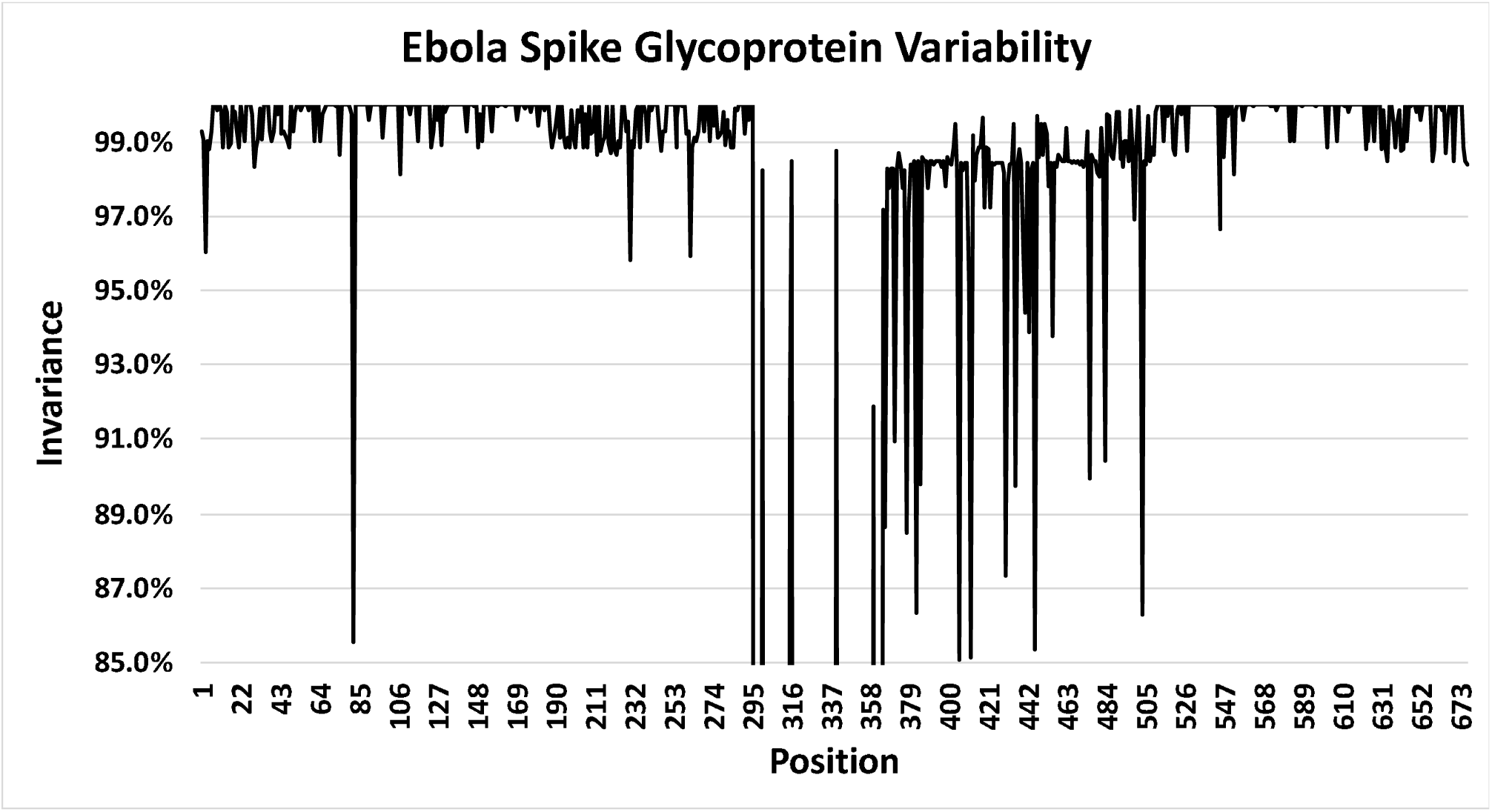
Ebola spike glycoprotein variability plot by protein residue position

**Figure 4.**
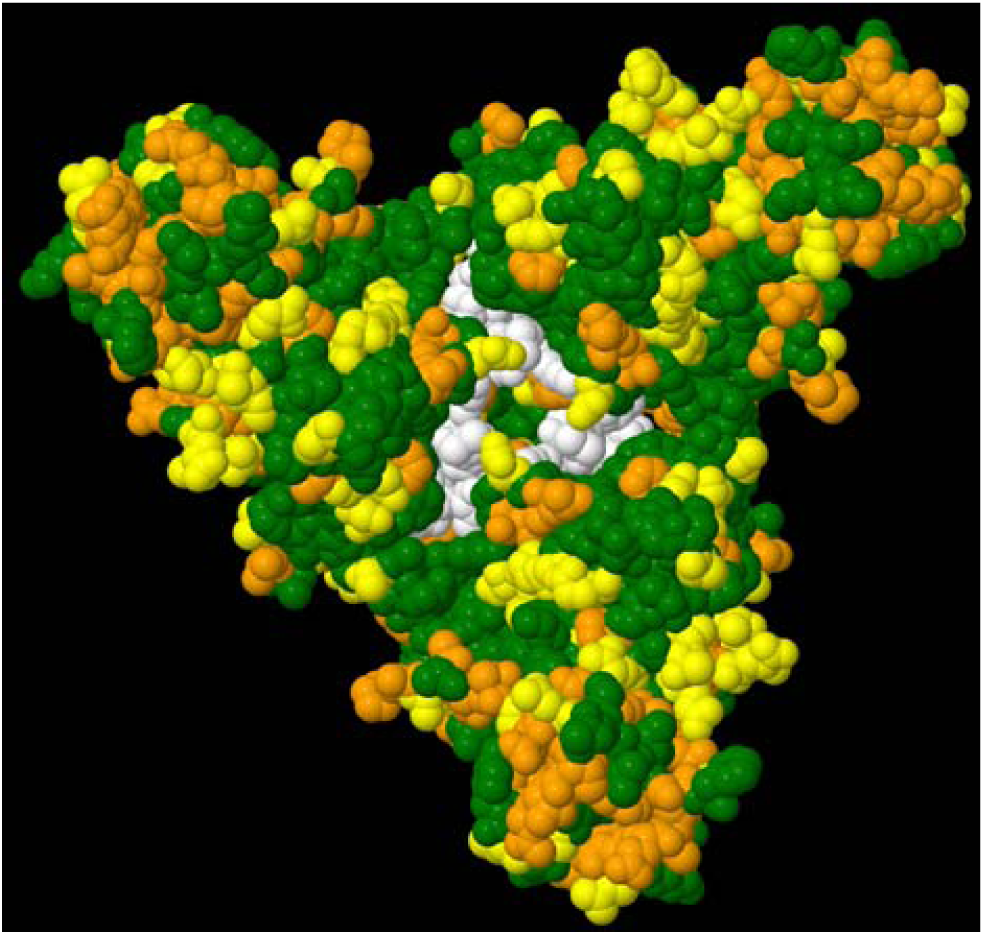
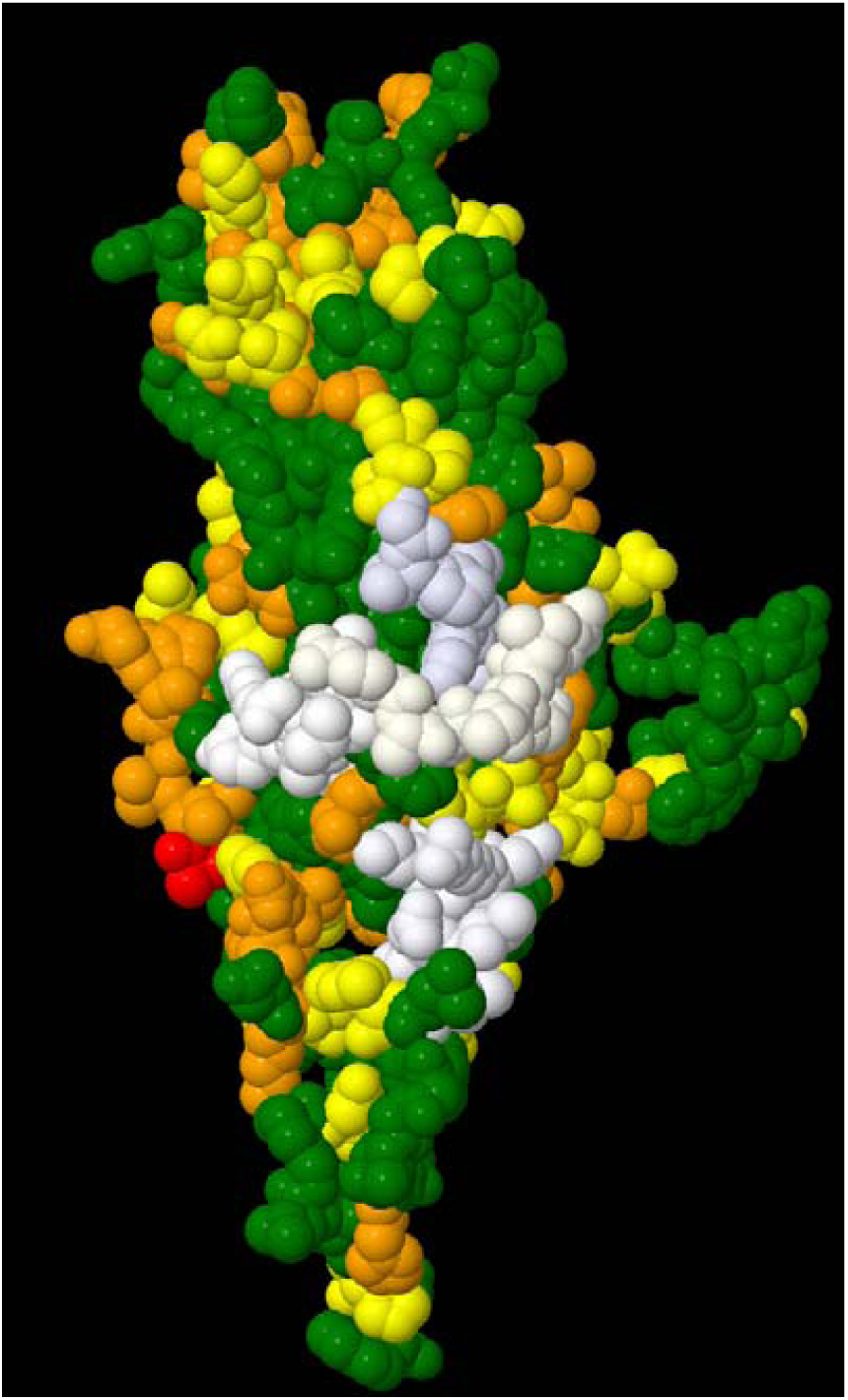
Ebola glycoprotein target peptides NETTQALQL, RATTELR, and NRKAIDFL. Residue color code used for 5KEL: white – target candidates, green – non-variable residues, yellow – up to 1% variability, orange – up to 5% variability, and red – up to 10% variability. (a) Glycoprotein trimer with buried interface residues in white (b) Glycoprotein monomer with exposed surface interface residues: target linear peptides NETTQALQL (color white), RATTELR (color ivory), and NRKAIDFL (ghostwhite)

**Figure 5.**
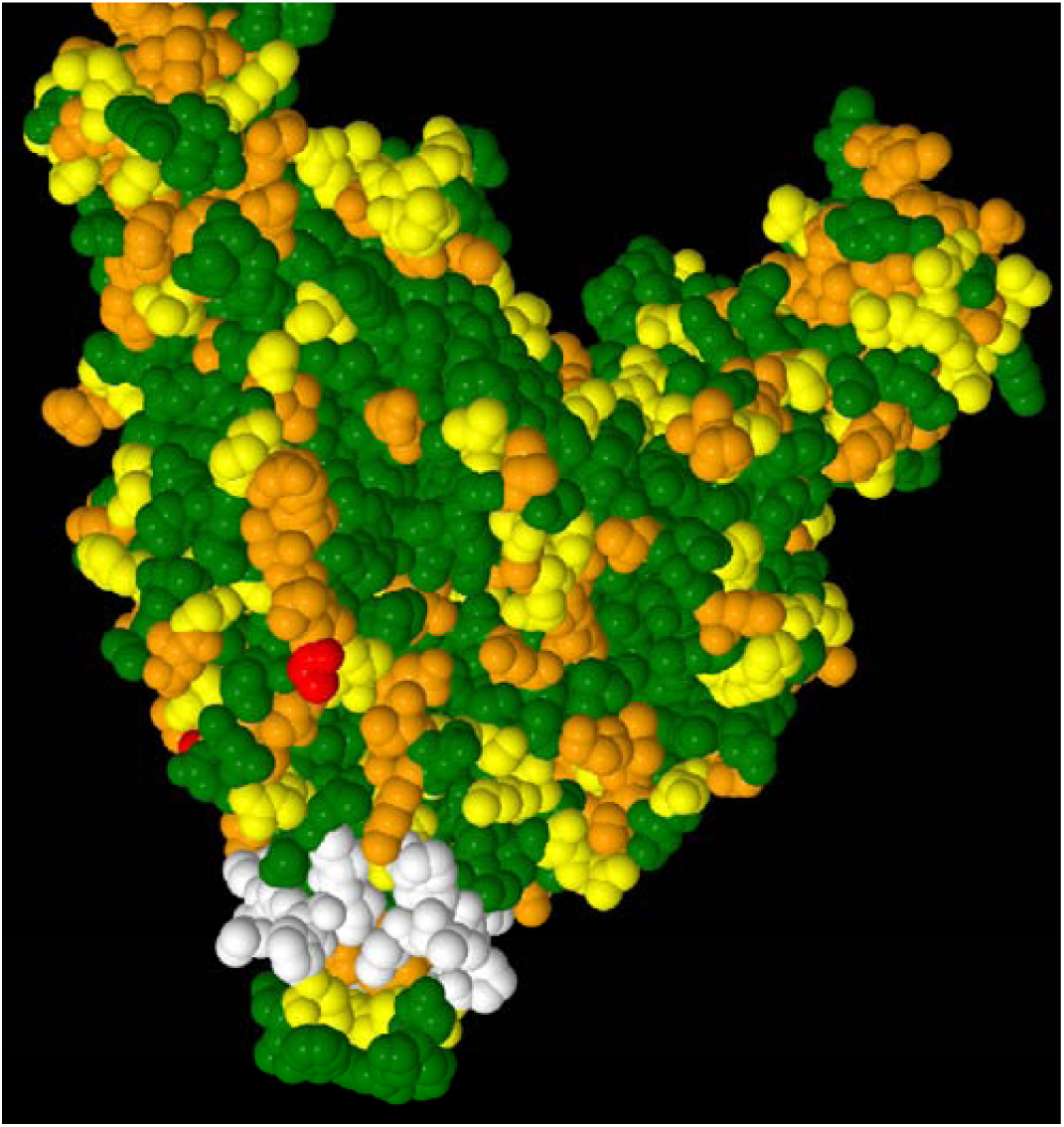
Ebola glycoprotein epitope ILGPDCCIEP

**Figure 6.**
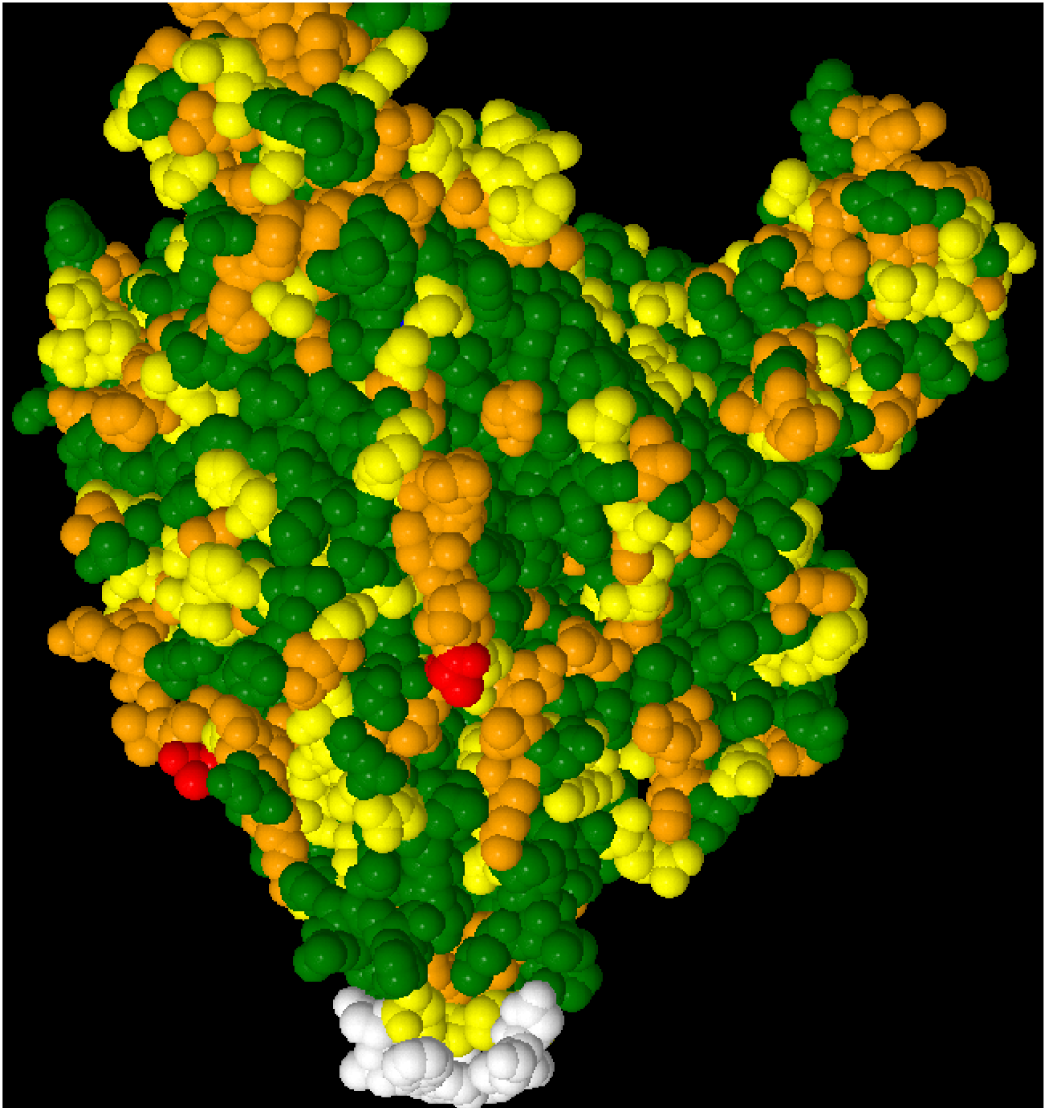
Ebola glycoprotein epitope DWTKNITD

**Figure 7.**
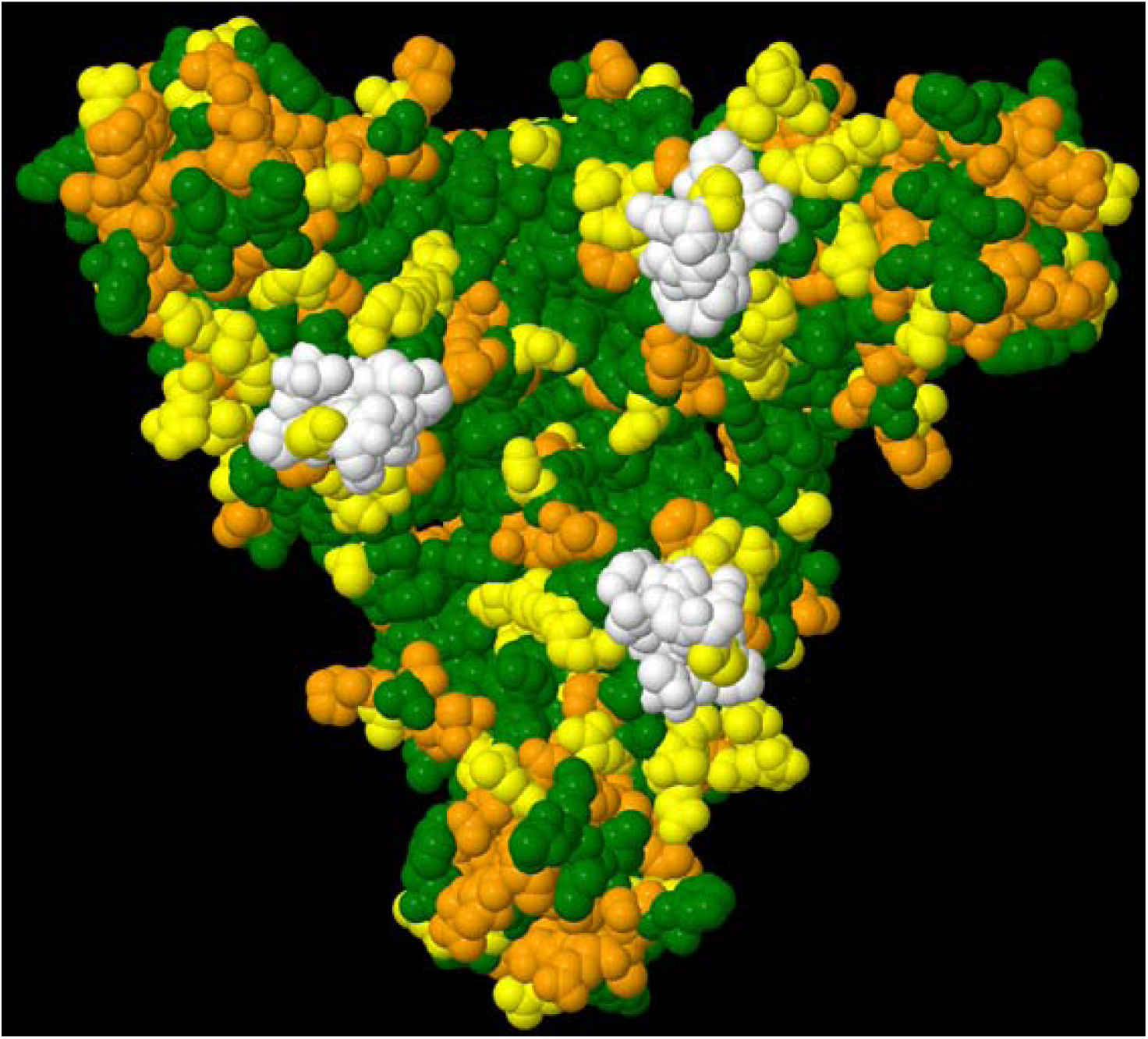
Ebola glycoprotein exposed surface target residues in glycan cap 2: K_114_D_117_S_119_I_120_L_122_G_143_T_144_P_146_

## Discussion

The variability analysis methodology introduced for targeting influenza hemagglutinin and neuraminidase[12] has been applied to the Ebola glycoprotein. Ebola glycoprotein residues with no or low observed variability identified multiple candidate targets antibody epitopes (Table 1). Linear peptide targets with no variability are also targets for T-cell vaccination strategy, RNA targeting (Table 2), and aptamer targeting. A similar strategy was recently developed for targeting multiple MAGE-A family members simultaneously for cancer therapy[22]. These computational targets can be evaluated in parallel for rapidly developing countermeasures to viral and bacterial pathogens. The target peptides can be combined together into a DNA or mRNA vaccine that targets one or ideally multiple pathogen target proteins. These targets can also serve as synthetic miRNA/siRNA and aptamer anti-virus strategies. Traditional viral subcomponent nucleic acid[1] and the multimeric low variability linear peptides (T-cell response stimulation) serve as population vaccination approaches. The bnAb, anti-RNA, and aptamer targeting strategies serve as treatment approaches for infected individuals (Figure 1). Bounds *et al*.[23] identified three cluster sequences that overlap with the candidate targets TKRWGFR, NETTQALQ, and RATTELRT in a multi-epitope construct; this construct by itself did not provide protection to virus challenged mice. The candidate targets TKRWGFR and NETTQALQ were also identified by Yasmin & Nabi[24] as subsets of longer peptides. A larger B cell epitope was identified by Babirye *et al*.[25] that includes both candidate targets NETTQALQ and RATTELRT from Ebola virus disease (EVD) survivor sample. Ebola immune evasion strategies include the down regulation of type I interferon production, masking of viral epitopes with glycosylation, and the secretion of sGP to overwhelm the host’s adaptive humoral immunity. The four targets TKRWGFR, GVPPKVV, DGSECLP, and KDSILGTP would also be in the secreted glycoprotein (sGP); the remaining targets after residue 324 would not be affected by the secretion of sGP. The strategy to target the Ebola genome and RNA molecules with synthetic complimentary RNA molecules may not be effective due to the Ebola suppression of the interferon response to double stranded RNA molecules but may be effective against other viral pathogens. Table 3 summaries the candidate targets for each approach presented.

**Table 3.**
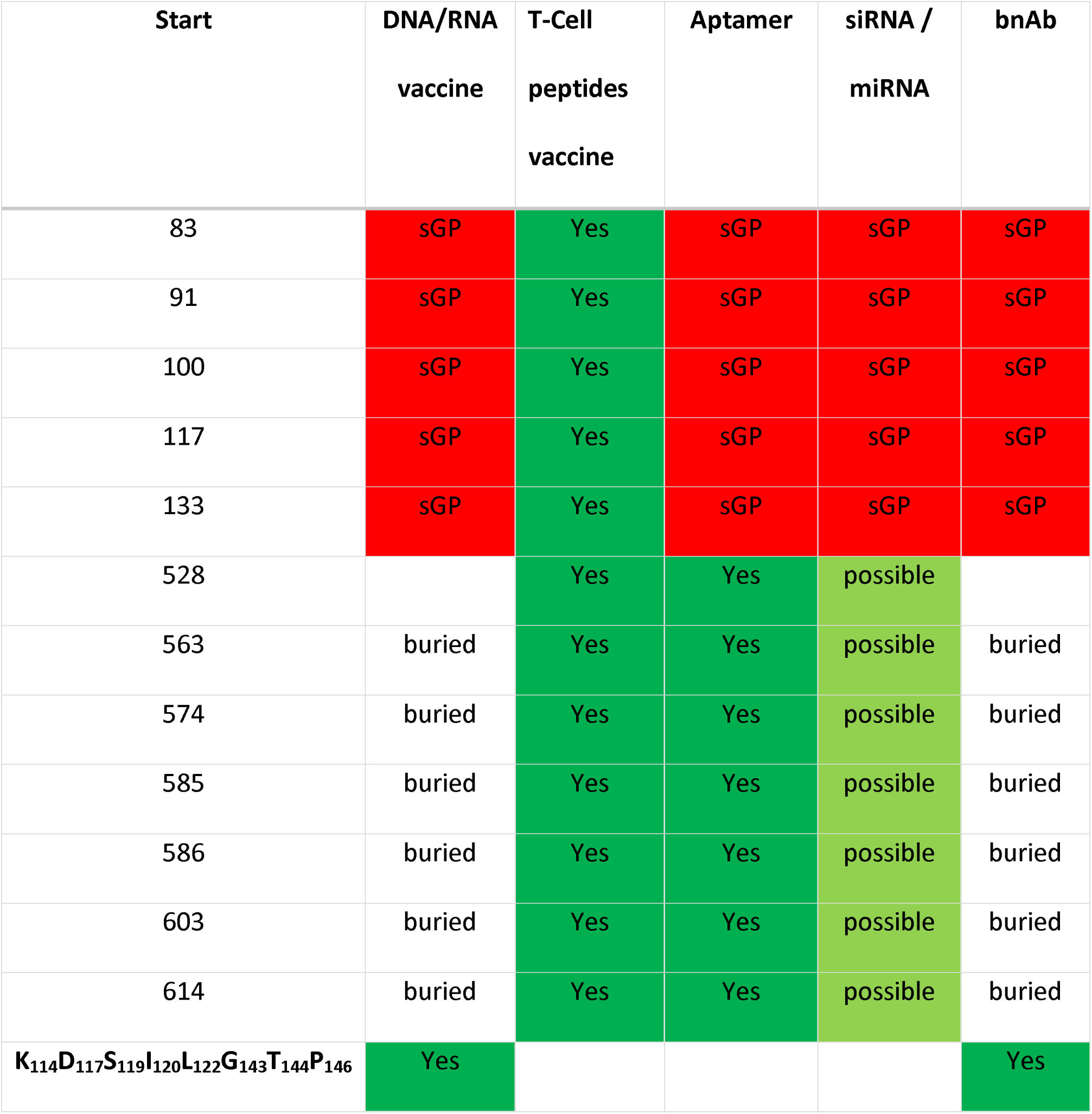
Ebola glycoprotein layered response targets; epitopes and RNA targets also contained in sGP are not considered good targets (red shading).

| Start | DNA/RNA<br>vaccine | T-Cell<br>peptides<br>vaccine | Aptamer | siRNA /<br>miRNA | bnAb |
| --- | --- | --- | --- | --- | --- |
| 83 | sGP | Yes | sGP | sGP | sGP |
| 91 | sGP | Yes | sGP | sGP | sGP |
| 100 | sGP | Yes | sGP | sGP | sGP |
| 117 | sGP | Yes | sGP | sGP | sGP |
| 133 | sGP | Yes | sGP | sGP | sGP |
| 528 |  | Yes | Yes | possible |  |
| 563 | buried | Yes | Yes | possible | buried |
| 574 | buried | Yes | Yes | possible | buried |
| 585 | buried | Yes | Yes | possible | buried |
| 586 | buried | Yes | Yes | possible | buried |
| 603 | buried | Yes | Yes | possible | buried |
| 614 | buried | Yes | Yes | possible | buried |
| **K<sub>114</sub>D<sub>117</sub>S<sub>119</sub>I<sub>120</sub>L<sub>122</sub>G<sub>143</sub>T<sub>144</sub>P<sub>146</sub>** | **Yes** |  |  |  | **Yes** |

The variability analysis methodology has only recently become practical for targeting pathogens. Computer scientists have classified multiple sequence alignment as NP-complete/NP-hard problem for which scalable solutions don’t exist[26]. Bioinformatics tools like MUSCLE[27], T-coffee[28], etc. either excessively over-gap alignments or simply fail when the number of sequences becomes too large. The recent development of tools like Dawn[12] and Clustal-Omega[13] enable high quality alignment of large numbers of protein sequences to enable variability analysis of pathogen proteins.

Newly discovered viruses may have limited numbers of available sequences. These available sequences can be aligned with evolutionarily related sequences and also modeled on available structures for these related sequences when no structures for newly discovered viruses are available. The Critical-Spacer[29] and Divergence models[30] for protein structure posit that the low variability residues are composed of shared functional residues and alignable core structures residues. Low variability residues selected from the alignment of multiple pathogen ortholog sequences can be leveraged as an alternative for having fewer sequences for the newly discovered virus.

## Summary

Variability analysis enables computational targeting of pathogens with multiple parallel approaches, including subcomponent and low variability peptide targets for stimulating B-cell and T-cell immune responses as part of vaccinations, bnAb targeting of low variability surface targets, anti-RNA targeting with synthetic miRNA/siRNA, and aptamer targeting of low variability surface targets. Parallel evaluation of these approaches can be evaluated *in vitro* and *in vivo* animal studies as part of a rapid medical countermeasures approach. Effectiveness of countermeasures may be enhanced with combination approaches of two or more of these countermeasures.

## Acknowledgements

DISTRIBUTION STATEMENT A. Approved for public release. Distribution is unlimited.

This material is based upon work supported under Air Force Contract No. FA8702-15-D-0001. Any opinions, findings, conclusions or recommendations expressed in this material are those of the author(s) and do not necessarily reflect the views of the U.S. Air Force.

The author would like to thank Irene Stapleford for artwork and Catherine Cabrera for feedback on this manuscript.

